# DNA Extraction Method Optimized for Nontuberculous Mycobacteria Long-Read Whole Genome Sequencing

**DOI:** 10.1101/470245

**Authors:** Jennifer M. Bouso, Paul J. Planet

**Affiliations:** Division of Pulmonary Medicine, Children’s Hospital of Philadelphia, Philadelphia, Pennsylvania, United States of America; Division of Infectious Diseases, Children’s Hospital of Philadelphia, Philadelphia, Pennsylvania, United States of America; Perelman School of Medicine, University of Pennsylvania, Philadelphia, Pennsylvania, United States of America; Sackler Institute for Comparative Genomics, American Museum of Natural History, New York, New York, United States of America

## Abstract

Nontuberculous mycobacteria (NTM) are a major cause of pulmonary and systemic disease in at-risk populations. Gaps in knowledge about transmission patterns, evolution, and pathogenicity during infection have prompted a recent surge in genomic NTM research. Increased availability and affordability of whole genome sequencing (WGS) techniques, including the advent of Oxford Nanopore Technologies, provide new opportunities to sequence complete NTM genomes at a fraction of the previous cost. However, extracting large quantities of pure genomic DNA is particularly challenging with NTM due to their slow growth and recalcitrant cell wall. Here we report a DNA extraction protocol that is optimized for long-read WGS of NTM, yielding large quantities of highly pure DNA. Our refined method was compared to 6 other methods with variations in timing of mechanical and enzymatic digestion, quantity of matrix material, and reagents used in extraction and precipitation. We also demonstrate the ability of our optimized protocol to produce sufficient DNA to yield near-complete NTM genome assemblies using Oxford Nanopore Technologies long-read sequencing.

## Introduction

The emergence of nontuberculous mycobacteria (NTM) infection in immunocompromised hosts, the elderly, patients with cystic fibrosis (CF), and patients with non-CF chronic lung disease (COPD, asthma, non-CF bronchiectasis) has prompted genomic investigations aimed at uncovering the determinants of pathogenicity, transmission, evolution, and adaptation (1-10). Recent studies of bacterial evolution and phylogenomics have been revolutionized by more available and affordable of whole genome sequencing (WGS) (11-15). Whole genome sequencing of NTM has begun to shed light on taxonomic conundrums, transmissibility, and global evolution (16-24). However, the unique challenges of slow growth rates and inefficient DNA extraction have impeded rigorous genomic investigation of NTM.

Over recent years, the vast majority of genomic analysis has relied on short-read, shot-gun sequencing (125-500 base pairs), which can deliver exceptional accuracy, but rarely produces closed genomes. Indeed, less than 10% of available microbial genomes are complete (25). Fragmented assemblies are problematic because they may unlink gene clusters, fail to resolve repetitive and G+C rich regions, neglect insertion and deletion elements (indels), and overlook recombination (26-29).

Long-read sequencing promises an enhanced ability to complete bacterial genomes. The most commonly available techniques for long-read sequencing are the Single Molecule Real-Time (SMRT) technology by Pacific Biosciences^®^ (PacBio) and the newer Oxford Nanopore Technologies (ONT) (14, 27, 30). Unlike most short-read sequencing methods, which require only very small amounts of DNA (as low as 1 ng), long-read platforms require high quantities of very pure DNA for acceptable processing (Table 1). DNA purity and integrity (*i.e*., length or molecular weight [MW]) is not only essential for functionality of the sequencer, but also is directly related to the quality of downstream bioinformatic analyses, as the DNA MW places a natural upper bound on the potential read length.

**Table 1.** DNA specifications by long-read WGS platform.

|  | PacBio | MinION |
| --- | --- | --- |
| Minimum Input DNA | 5 µg | 400 ng |
| 260/280 | 1.6-1.8 | 1.6-1.8 |
| 260/230 | 2.0 | 2.0 |
| Expected Library Insert Size | 500bp – 20 Kb | unlimited |
| Expected DNA read length | up to 500 Kb | unlimited |
| Expected GB per flow cell | 20 GB | 10 – 20 GB |

Extracting large quantities of intact, pure genomic DNA is exceptionally challenging with NTM due to their hardy, lipid-laden mycobacterial cell wall. Standard extraction techniques do not yield sufficient quantities of DNA for WGS while overly vigorous techniques shear DNA into suboptimal MWs for long-read sequencing. Previously, Käser et al. in 2010 published a mycobacterial-specific DNA extraction protocol, which is commonly used in the NTM research community (31, 32). In our experience, this technique was unable to yield DNA that was of sufficient quality for ONT MinION sequencing. We developed an optimized protocol over the course of performing over 100 NTM DNA extractions, using components of several extraction techniques (31, 33-36). Our improved method is characterized by early bead-beating (prior to enzymatic digestion) in high concentrations of sodium dodecyl sulfate (SDS) followed by gentle extraction and precipitation focused on DNA purification and protection of long strands of DNA. The goal of our protocol is to extract high MW, pure DNA for use in long-read WGS. Here, we demonstrate its superiority to 6 variations in methodologic design and validate its capacity for producing near-complete genome assemblies with the ONT MinION sequencer.

## Materials and Methods

### Bacterial growth

Clinical isolates of *M. avium* complex were grown from frozen stocks to Löwenstein–Jensen slants and sub-cultured to Middlebrook 7H11 plates. Single colonies from 7H11 plates were inoculated in Middlebrook 7H9 broth supplemented with 10% OADC and incubated statically at 37°C for 2 weeks. Bacterial cultures were pelleted (4500 rpm x 10 min) and stored at −20°C until time of extraction.

### DNA extraction

The following extraction protocol described is “Method 5.” Alternate methods are described in Table 2. Method 3 corresponds to the protocol by Käser et al (32). The comprehensive protocol with thorough descriptions of each step and reagent recipes is provided in Supplemental Figure 1.

**Table 2.** Differentiation of Tested Methods by Variable.

|  | Method 1 | Method 2 | Method 3 | Method 4 | Method 5 | Method 6 | Method 7 |
| --- | --- | --- | --- | --- | --- | --- | --- |
| Early vs. Late bead-beating* | Early | Late | Late | Early | Early | Early | Early |
| Bead Quantity | 150 mg | 150 mg | 150 mg | 75 mg | 150 mg | 150 mg | 150 mg |
| Phenol vs. No Phenol† | No phenol | No phenol | Phenol | No phenol | Phenol | No Phenol | No phenol |
| Precipitation Temp/ Reagent‡ | RT/2-Prop | RT/2-Prop | Cold ETOH | RT/2-Prop | RT/2-Prop | Cold ETOH | RT/2-Prop |
| Precipitation Salt§ | NaOAc | NaOAc | NaOAc | NaOAc | NaCl | NaOAc | NaOAc |
| Number of washes | 3 | 3 | 3 | 3 | 3 | 3 | 5 |

\* "Early" bead-beating refers to the timing *prior* to enzymatic digestion; "Late" bead-beating refers to timing *after* enzymatic digestion.
All Early bead-beating was done in high SDS concentration, see Supplemental Figure 1.
† DNA extractions in "Phenol" were extracted as described in the Materials and Methods with phenol:chloroform:isoamyl alcohol (25:24:1, Tris-saturated, pH 8.0); extractions in "No phenol" were extracted using chloroform:isoamyl alcohol (24:1, Tris-saturated).
‡ Precipitation reagent was either RT 2-Prop (room temperature isopropanol) or Cold ETOH (100% ethanol chilled at -20°C).
§ Precipitation salt was either 3 M sodium acetate (pH 5.2) or 5 M NaCl.

#### Sample Preparation

Bacterial pellets were resuspended and washed in 350 µL of 1X phosphate-buffered saline (PBS) using 2 mL microcentrifuge tubes. Due to variability in starting mass between bacterial isolate cultures, and for the purposes of comparing extraction methods, all weights were normalized after a 2^nd^ PBS wash and “washed weights” were recorded. The samples were heat-inactivated for 60 minutes at 95°C, pelleted, and supernatant discarded.

#### “Early” Mechanical Disruption in SDS followed by Enzymatic Digestion

Bacterial pellets were resuspended in 400 µL of lysis buffer and 100 µL of 20% SDS. Samples were homogenized with glass beads (4 × 30 second cycles at 3000 rpm, Fisher Scientific vortex mixer, MoBio adapter) (150 mg glass beads, 0.1-mm diameter, Research Products International). Subsequently, all vortexing was avoided. Cell walls were additionally lysed in lysozyme (final concentration 10 mg/mL) for 1 hour at 37°C.

Proteinase K (final concentration 200 µg/mL) was added and samples were incubated at 37°C for 90 minutes with mixing by turning end-over-end by hand every 30 minutes. The lysates were centrifuged (4500 rpm × 10 min followed by 14,000 rpm x 2 min) and supernatants transferred to 2 mL 5Prime Light Phase Lock Gel™ (PLG, QuantaBio) microcentrifuge tubes. Variables tested in the aforementioned digestion steps included enzymatic digestion prior to mechanical digestion (Methods 2, 3) and the amount of matrix material added (Method 4).

#### Phenol:Chloroform:isoamyl alcohol extraction

To extract DNA, 500 µL of phenol:chloroform:isoamyl alcohol (25:24:1, Tris-saturated, pH 8.0) was added to the PLG tubes. The tubes were rotated on a HulaMixer at 20 rpm for 20 minutes and then centrifuged (4500 rpm × 10 min). The DNA-containing aqueous layer was transferred to a new 2 mL microcentrifuge tube. Chloroform:isoamyl alcohol (24:1, Tris-saturated) without phenol was tested as a variable (Methods 1, 2, 4, 6, 7).

#### Isopropanol precipitation

For DNA precipitation, 1/10 volume of 5 M sodium chloride (~20-45 µL) and 1 volume of room temperature isopropanol (~200-450 µL) was added to the samples. The samples were incubated at room temperature overnight. The samples were then centrifuged (14,000 rpm × 30 min at 22°C, to avoid heating), washed with 700 µL 70% ethanol (14,000 rpm × 10 min at 22°C), and the supernatant carefully discarded, with repeat of washing steps 3 times. The samples were air-dried at room temperature with lids open for 15 minutes, resuspended in 100 µL of Tris-Cl Elution Buffer (Qiagen), and eluted overnight on a nutator (24 rpm fixed speed, Fisherbrand™). DNA was stored at 4°C for 3-4 days prior to quality assessment. Variables tested during precipitation include use of cold 100% ethanol (Methods 3, 6) and use of an alternative salt (Methods 1-4, 6, 7).

#### Quality measures

DNA was heated to 65°C × 1 hour prior to quality assessment. DNA purity was assessed with NanoDrop 2000 UV-Vis Spectrophotometer (260/280, 260/230) and concentrations measured with Qubit^®^ 2.0 Fluorometer (dsDNA BR Assay). Gel electrophoresis (0.6% agarose ethidium bromide gel, 40V × 2 hours) estimated molecular weights and shearing. One-way ANOVA with post-hoc Tukey’s multiple comparison test in Prism 7.0d for Mac OS × (GraphPad Software, La Jolla California USA, www.graphpad.com) was used to determine significance differences.

#### Whole genome sequencing

Two representative samples of *M. avium* subsp. *hominissuis* (CHOP101034 and CHOP101174) were prepared for WGS with library preparations of Nextera XT (Illumina) and Rapid Barcoding Kit (ONT), and libraries were sequenced on their respective platforms of Illumina HiSeq 2500 and ONT MinION sequencer (FLO-MIN107). Read qualities were assessed with FastQC and MultiQC (37, 38). Genome assemblies were constructed with short-reads only, long-reads only, and with a combination of short and long-reads (hybrid) with error correction. Raw reads were trimmed and demultiplexed with Trim Galore (39, 40), *de novo* assembled with Unicyler (41), and assembly graphs generated by Bandage (42). Long-read and hybrid assemblies were additionally polished and circularized with Circlator (43). Assembly quality control measurements were assessed with QUAST (44).

## Results

Initial bacterial pellets averaged a normalized “washed weight” of 26.4 mg. With the exception of Method 6, all methods tested produced sufficient total DNA quantity and concentration (Figure 1a, 1b). Methods 1 and 3 produced the highest total amounts of DNA with 12.45 and 11.43 µg of DNA, respectively (Table 3). All methods except Method 6 gave sufficient 260/280, indicating low protein contamination overall (Table 3, Figure 2a). Method 3 and 5 produced the highest 260/280 measurements, which were significantly higher than other methods. Only Method 5 produced sufficient 260/230 for use with long-read sequencers without the need for any clean-up steps (Table 3, Figure 2b). Despite apparent variation in DNA quantity, all methods produced high MW DNA as evidenced on an ethidium bromide gel, indicating preservation of long reads of genomic DNA (Figure 3).

**Table 3.**
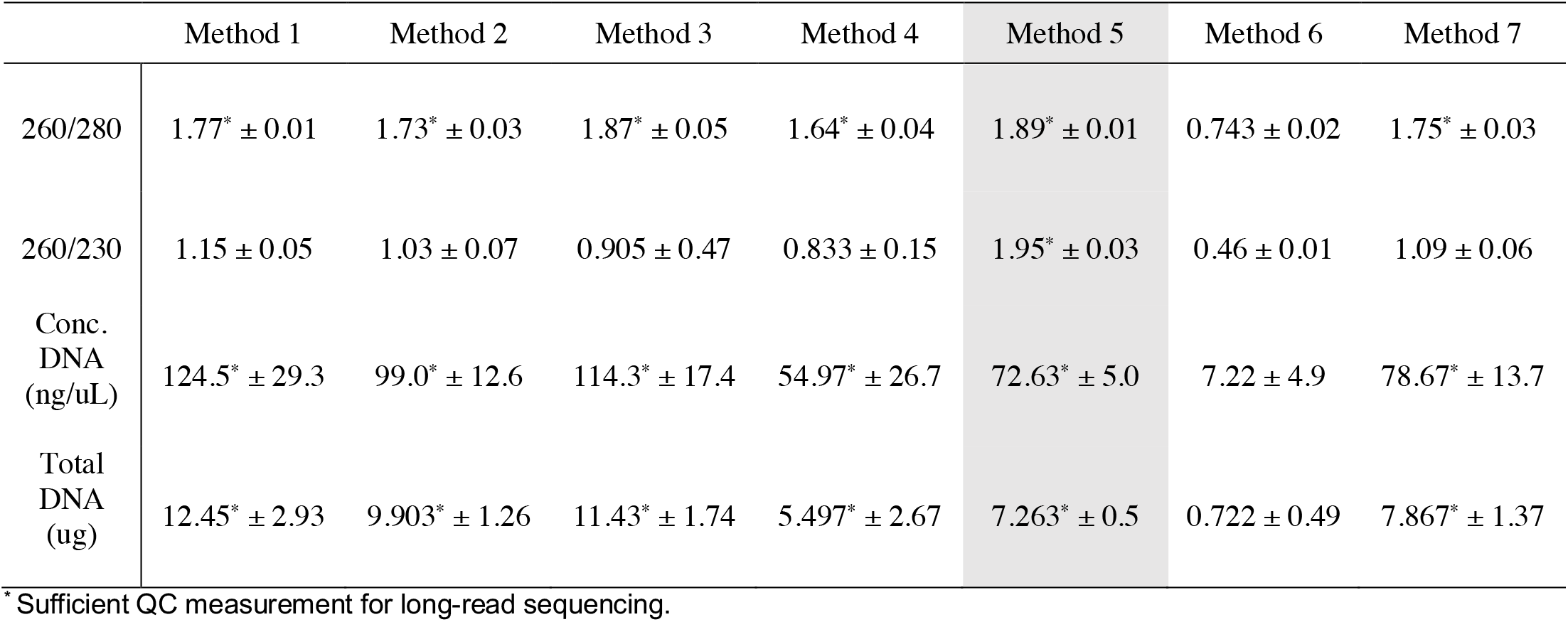
Average Quality and Quantity Measurements by Method. All reported values are averages of extractions performed in triplicate with ± standard deviation. The highlighted method achieved sufficient QC on all measurements.

**Table 4.** Quality Statistics of Isolates Sequenced for Long-Read Hybrid Assembly. Statistics were generated with FastQC and MultiQC (37, 38).

|  |  | Total reads | GB Data | Avg. read length (bp) | Longest read (bp) | %GC | Coverage* | Mean Phred score | % Error Probability† |
| --- | --- | --- | --- | --- | --- | --- | --- | --- | --- |
| Illumina | CHOP101034 | 2,521,793 | 1.6 | 126 | 126 | 65% | 122x | 37 | 0.01995% |
|  | CHOP101174 | 2,348,600 | 1.5 | 126 | 126 | 67% | 113x | 37 | 0.01995% |
| ONT | CHOP101034 | 545,494 | 1.3 | 1,139 | 34,615 | 66% | 119x | 13 | 5.0119% |
|  | CHOP101174 | 875,670 | 2.0 | 1,122 | 36,523 | 66% | 188x | 13 | 5.0119% |

\* Coverage calculated by $C = LN/G$ , where L=average read length, N=number of reads, G=genome size of 5.2 Mb.
† Error probability percentage is a function of Mean Phred score, where probability $P\% = 100 \cdot 10^{(-Phred/10)}$ .

**Figure 1.**
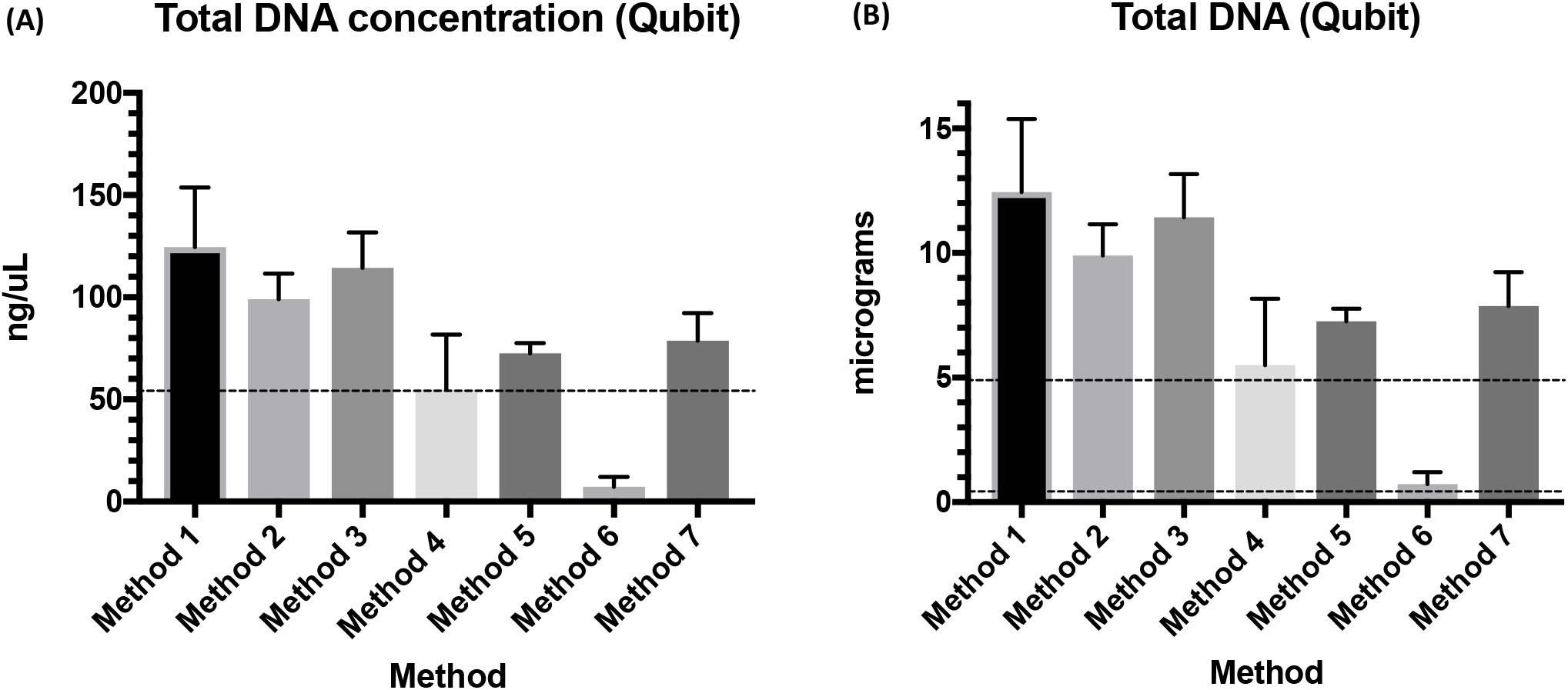
Quantification of DNA by Method. (A) DNA concentration as measured by Qubit fluorometry with dotted line indicating the minimum required input DNA concentration for PacBio (PB) and Oxford Nanopore Technologies MinION (ONT). (B) Total DNA as measured by Qubit fluorometry with dotted lines indicating minimum required input DNA for PB and ONT sequencing.

**Figure 2.**
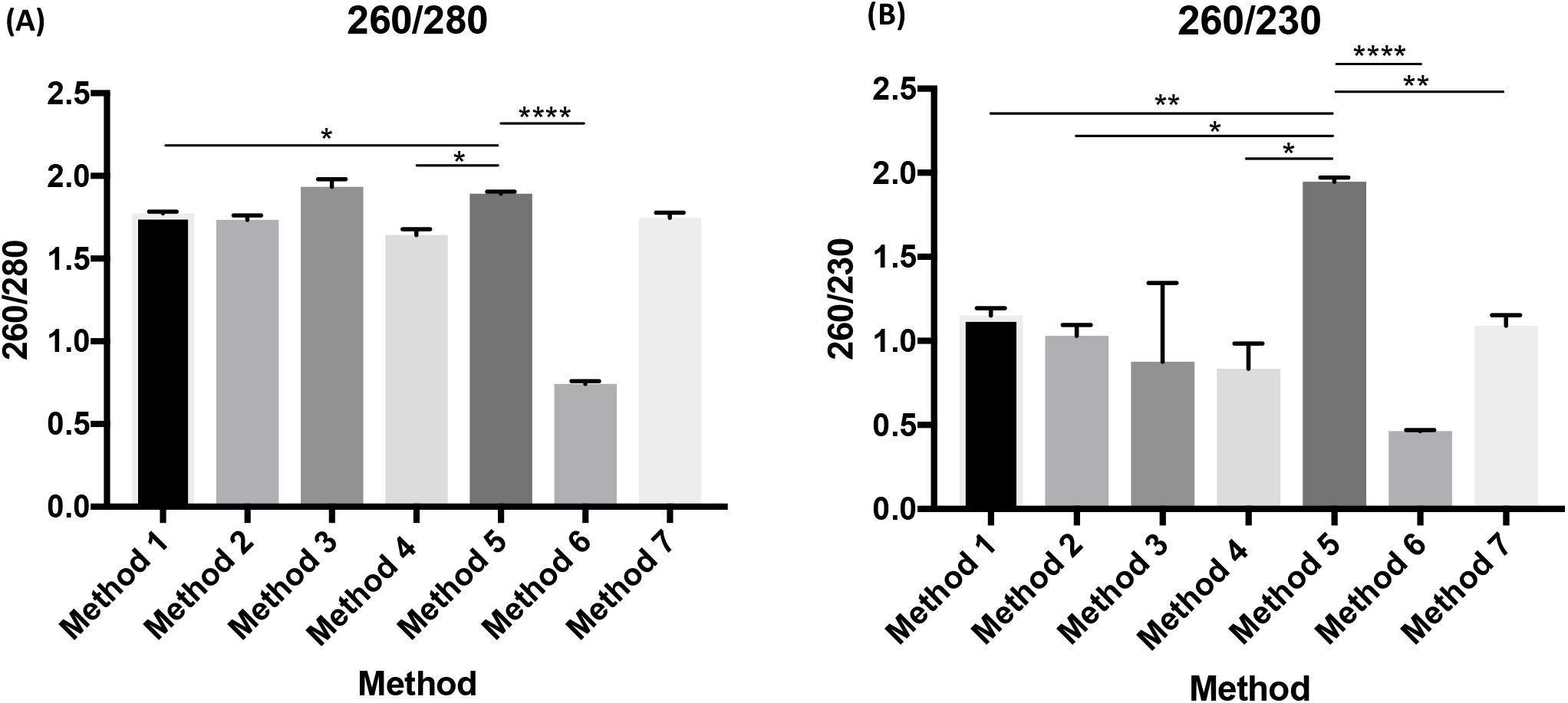
Quality and Quantity Measurements by Method. All values are presented as averages of each tested method in triplicate with error bars indicating standard deviation. Significance was computed with one-way ANOVA with post-hoc Tukey’s multiple comparison test. For clarity, significance bars were only depicted for comparisons against Method 5. (A) 260/280 by method and (B) 260/230 by method. Significance depicted by p-values as follows: 0.0332 (*), 0.0021 (**), 0.0002 (***), <0.0001 (****).

**Figure 3.**
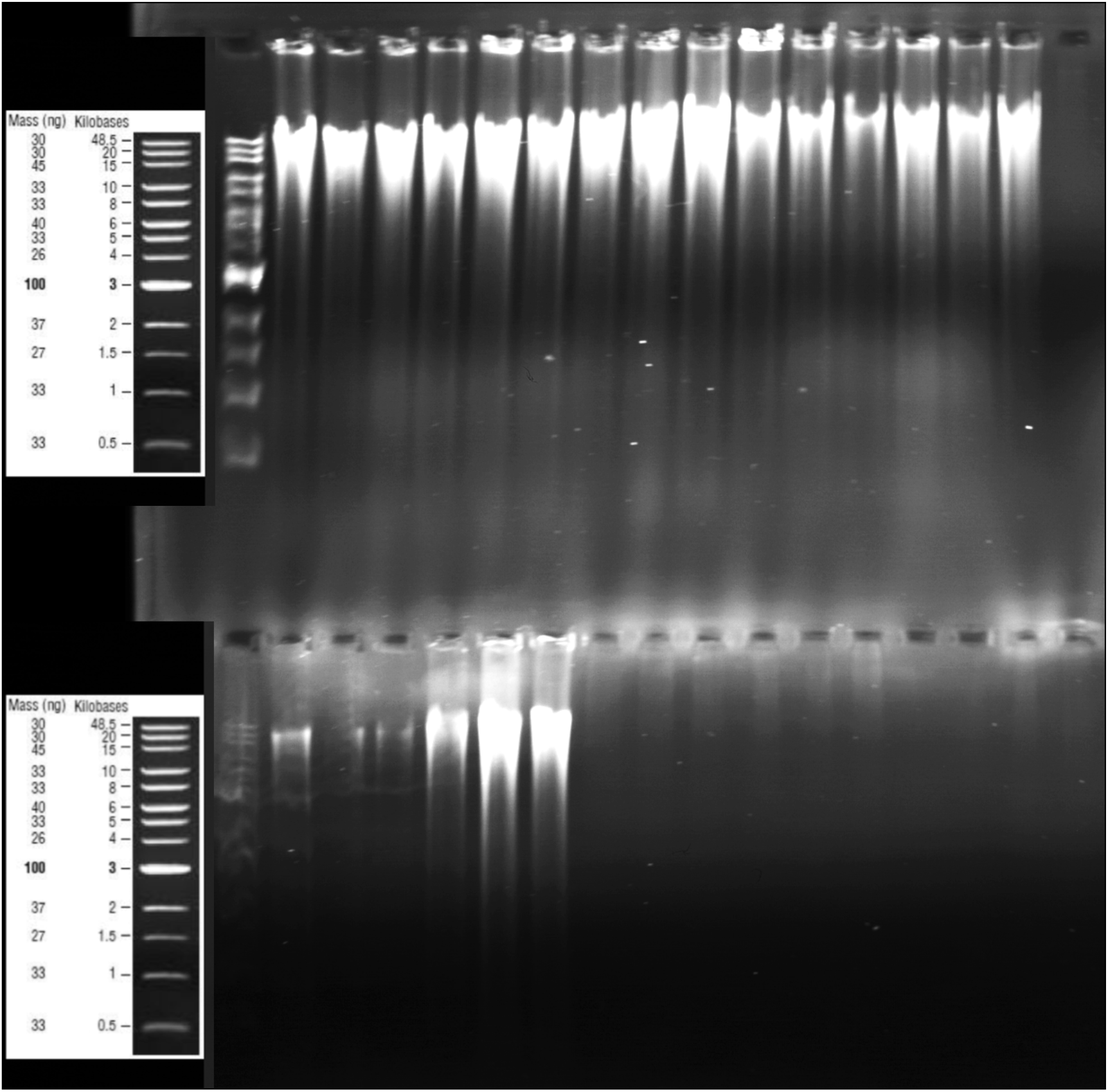
Gel Electrophoresis of gDNA. Methods 1-7 in triplicate (left to right) on a 0.6% EtBr gel demonstrating high MW genomic DNA.

Representative samples of DNA from single colony extractions (CHOP101034 and CHOP101174) were sequenced by both Illumina NGS and ONT MinION with quality control statistics of reads listed in Table 3. Remarkably, ONT reads averaged over 1000 bp without size selection with longest reads of approximately 35,000 bp. Such high read lengths supplied higher coverage for ONT reads despite lower base calling accuracy. In this regard, Illumina and ONT read qualities generated expected results. Comparison of mean Phred scores of Illumina reads (37) and ONT reads (13) demonstrated a lower probability of a miscalled base at any given position with Illumina sequencing (0.02% versus 5.01%).

Quality control statistics of assembled genomes computed in QUAST and assembly graphs generated by Bandage are featured in Figure 4 (42, 44). Short-read only assemblies were considerably more fragmented with an average of 90 contigs compared to the long-reads only and hybrid assemblies, which averaged 13 and 7 contigs, respectively. The best assembly (CHOP101174, hybrid) was 5.15 Mb in length with 5 contigs and an N50 of 3.0 Mb. Notably, ONT-only assemblies were overall similar to hybrid assemblies by basic quality statistic measures, but have significantly more errors based on Phred scores of input reads (Table 3).

**Figure 4.**
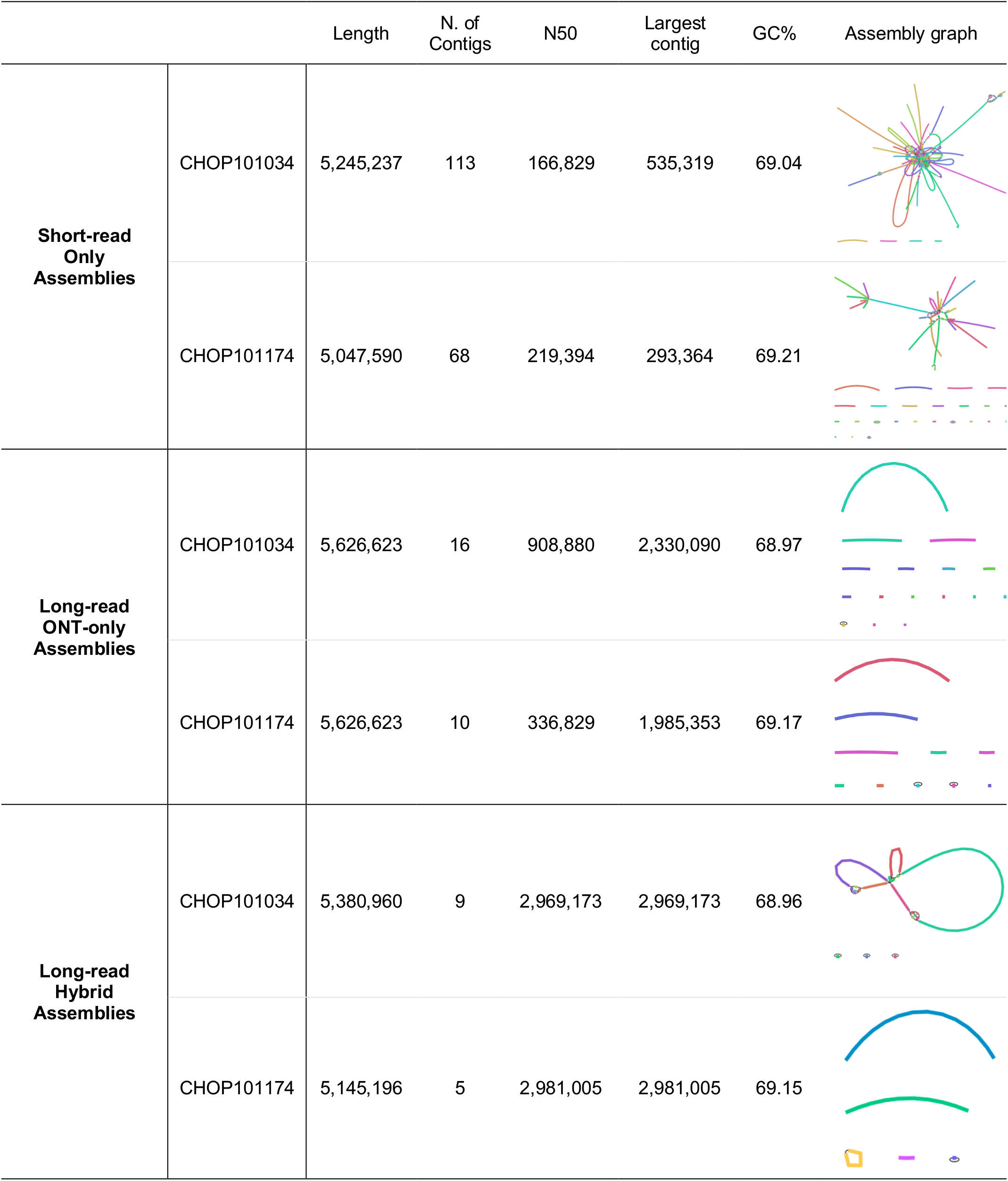
Assembly Statistics and Graphs. All assemblies were *de novo* assembled in Unicycler (41) with hybrid assemblies circularized with Circlator (43). Assembly statistics by QUAST (44) demonstrate considerably more complete assemblies with utilization of ONT long-reads and ssembly graphs generated by Bandage (42) provide visualization of more complete assemblies with long-read based assemblies.

## Discussion

Our protocol was used to supply sufficient input DNA for long-read WGS with the ONT MinION sequencer. To demonstrate direct comparisons to alternative methods, we completed DNA extraction with 6 variations of methodology with normalized starting DNA for head-to-head comparisons. Method 5 demonstrated superiority as the only method to provide appropriate quality control measurements without requiring any clean-up steps. Method 5 was characterized by early bead-beating in high-SDS concentration, gentle phenol-based extraction, and room temperature isopropanol precipitation. While Method 5 was the only method to use NaCl as the precipitation salt, later direct comparisons of NaCl versus NaOAc alone did not demonstrate any superiority of NaCl. Thus, while either salt is appropriate, we recommend NaCl over NaOAc because it does not require pH titration. In addition to the variables presented in this manuscript, we also trialed an alternative buffer, higher concentrations of lysozyme and proteinase K, variable starting weights of bacterial pellets, extraction without bead-beating, and bead-beating with and without SDS,

In comparison to a widely-used method (Method 3, Käser et al.), we noted improvements in the purity of DNA (260/280 and 260/230) with modifications of the composition of lysis buffer (See Supplemental Figure 1), the timing of bead beating (early vs. late), the use of Phase Lock Gel™ tubes, and precipitation in room temperature isopropanol as opposed to cold 100% ethanol. Others have shown improved DNA purity with isopropanol extractions compared to cold ethanol extractions with less salt carry-over, albeit at the expense of reduced DNA yields (31, 32). While Method 1 and 3 gave more total DNA, neither reached a suitable 260/230. Thus, total DNA yield may need to be sacrificed in order to achieve high DNA purity.

The trademark of the mycobacterial cell wall is its heavily lipophilic exterior. In addition, mycobacterial peptidoglycans are characterized by an oxidation modification rendering lysozyme less effective at cleaving the β(1,4) linkages between N-acetylmuramic acid and N-acetyl-D-glucosamine residues (45). Thus, it is no surprise that mechanical digestion is necessary for DNA extraction. We reason that *early* mechanical digestion allows the exterior mycolic acid cell wall and peptidoglycan layer to be broken down first, with subsequent enzymatic disruption with lysozyme and proteinase K to digest the remainder of the cell wall and expose the cellular contents. In our preliminary trials, early mechanical digestion demonstrated superiority to late mechanical digestion. Although not seen in the head-to-head comparisons presented here, we have noted increased shearing with late mechanical digestion, resulting in homogenously distributed smears of lower MW DNA on gel electrophoresis. In addition, we found that early addition of high concentrations of SDS during early beat-beating was also independently superior to early bead-beating without SDS (data not shown). The detergent properties of SDS likely assist with mechanical lysis and may additionally protect exposed DNA from degradation.

There are several reasons why long-read sequences are important for the interpretation of genomic evolution and phylogeny. Studies aimed at understanding patterns of transmission and intra-host adaptation must pay specific attention to unique genomic characteristics, such as mosaicism (46), recombination tracks (19), and large repeat regions (28, 29, 47), which can all function to confound phylogenetic inference. Current short-read techniques cannot obtain the contig sizes necessary to detect these genomic features. Complete WGSs also allow for optimal reference sequence generation for comparison of clonal relatives because they contain a more complete picture of genome content and organization and may detect genomic changes that would be missed otherwise. Thus, with the exception broad phylogenetic estimations, comprehensive variation-based analyses warrant supplementation with long-read assemblies.

Undoubtedly, long-read assembled genomes are the way of the future. As technology improves, assembly construction will be less and less reliant on short-read sequencing. However, we will remain at the mercy of the cell wall, and continue to be faced with the delicate challenge of mining unscathed DNA from a distinctly robust substrate. Here, we presented a finely-tuned extraction method designed for preparation of highly purified DNA to be used for long-read sequencing and demonstrated the ability to produce complete (or near complete) genome assemblies.

## Acknowledgements

This research was supported by the Cystic Fibrosis Foundation (BOUSO17BO). The funders had no role in study design, data collection and interpretation, or the decision to submit the work for publication.

